# Measuring the density and viscosity of culture media for optimized computational fluid dynamics analysis of *in vitro* devices

**DOI:** 10.1101/2020.08.25.266221

**Authors:** Christine Poon

## Abstract

Culture medium is frequently modelled as water in computational fluid dynamics (CFD) analysis of *in vitro* culture systems involving flow, such as bioreactors and organ-on-chips. However, culture medium can be expected to have different properties to water due to its higher solute content. Furthermore, cellular activities such as metabolism and secretion of ECM proteins alter the composition of culture medium and therefore its properties during culture. As these properties directly determine the hydromechanical stimuli exerted on cells *in vitro*, these, along with any changes during culture must be known for CFD modelling accuracy and meaningful interpretation of cellular responses. In this study, the density and dynamic viscosity of DMEM and RPMI-1640 media supplemented with typical concentrations of foetal bovine serum (0, 5, 10 and 20% v/v) were measured to serve as a reference for computational design analysis. Any changes in the properties of medium during culture were also investigated with NCI-H460 and HN6 cell lines. The density and dynamic viscosity of the media increased proportional to the % volume of added foetal bovine serum (FBS). Importantly, the viscosity of 5% FBS-supplemented RPMI-1640 was found to increase significantly after 3 days of culture of NCI-H460 and HN6 cell lines, with distinct differences between magnitude of change for each cell line. Finally, these experimentally-derived values were applied in CFD analysis of a simple microfluidic device, which demonstrated clear differences in maximum wall shear stress and pressure between fluid models. Overall, these results highlight the importance of characterizing model-specific properties for CFD design analysis of cell culture systems.

## 1. Introduction

Liquid culture media have been integral to the culture of mammalian cells since the inception of *in vitro* techniques and are composed of a mix of water, essential nutrients, vitamins and factors that support and regulate the growth of cells [1]. Other than the substrate, the culture medium is the other immediate environment that cells contact *in vitro*. In culture devices that involve flow such as bioreactors [2-7] and microfluidic organ-on-chips [8-11], culture fluid is the primary material through which mechanical loading is exerted on cells. However, significantly less attention has been paid to the physical properties of culture medium and how these may affect flow mechanics within these systems compared to considerations such as substrate geometry, tubing dimensions and flow rate [12]. Given the known sensitivity of cells to shear stimuli and particularly where the goal is to study the effects of physiological or pathological shear [13-17], it is essential to characterize precise flow properties and hydrodynamic regimes in order to (i) deliver known and controlled mechanical stimuli to cells, and (ii) correlate these to cellular responses.

Such analysis can be carried out via computational fluid dynamics (CFD) modelling, an effective method for quantitatively determining fluid flow phenomena that may be too complex or challenging to measure across a whole system or on cell-relevant (micron) scales by experimental means, e.g. by particle imaging velocimetry (PIV) [18, 19], laser doppler velocimetry [20-22], physical probes or sensors. CFD simulations are routinely conducted as part of the design process of fluid systems in engineering and have been increasingly implemented by tissue engineers and the microfluidics community to evaluate flow characteristics within device designs prior to prototyping [23-29]. With appropriate model set up, CFD analyses can provide accurate approximations of flow fields, velocity and stress profiles within tissue culture systems for design refinement [30, 31] and facilitate more precise study of the biological effects of fluid shear on cells.

The basic principle of CFD modelling is to resolve a set of governing equations that mathematically describe flow behaviours and properties across the discretized geometry (mesh) of a fluid system. Navier-Stokes and continuity equations (Equations 1 and 2) are the most applicable to the modelling of incompressible, Newtonian, water-like fluids, which aqueous culture media are typically considered to be for practical purposes. These equations are respectively based on conservation of momentum and mass and together, describe the space-time evolution of a velocity field in terms of convection, pressure, viscous drag and gravity. The equations are as follows:

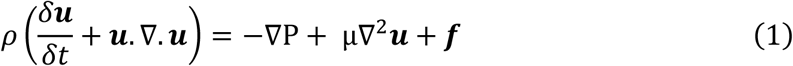

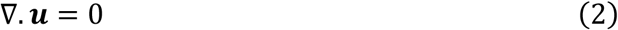

where ***u*** (m/s) denotes the fluid velocity vector, ∇ is the divergence operator, ρ (kg/m^3^) is the density and µ (Pa.s) is the dynamic viscosity of the fluid respectively, P (Pa) is pressure and **f** (m/s^2^) is gravity **g** where **f** = ρ**g** and/or an external acceleration field

From the Navier-Stokes equation (Equation 1), it can be seen that flow behaviour is directly determined by two key fluid properties-density and dynamic viscosity. CFD studies of *in vitro* systems typically model culture media as water at room temperature, where density and viscosity is assumed to be anywhere between 998-1000 kg/m^3^ and 0.8-1 mPa.s respectively [31-35]. A second assumption is that these fluid properties are constant [24, 36, 37], which is only applicable for continuous flow or single throughput systems where cells constantly receive fresh medium. Culture media can be expected to denser and more viscous than water due to its higher solute content (sugars, inorganic salts, sera proteins). Furthermore, these properties intuitively change over the course of an experiment as cells metabolize nutrients, growth factors and amino acids, secrete extracellular matrix proteins and excrete metabolic by-products [38], which changes the solute content and therefore the density, viscosity and hydrodynamics of the media (Figure 1). The degree and rate of change is a function of cell type, initial number of seeded cells, proliferation rate, the type and quantities of secreted proteins and other experimental factors such as the initial volume of medium and the rates of medium recirculation and replenishment.

**Figure 1.**
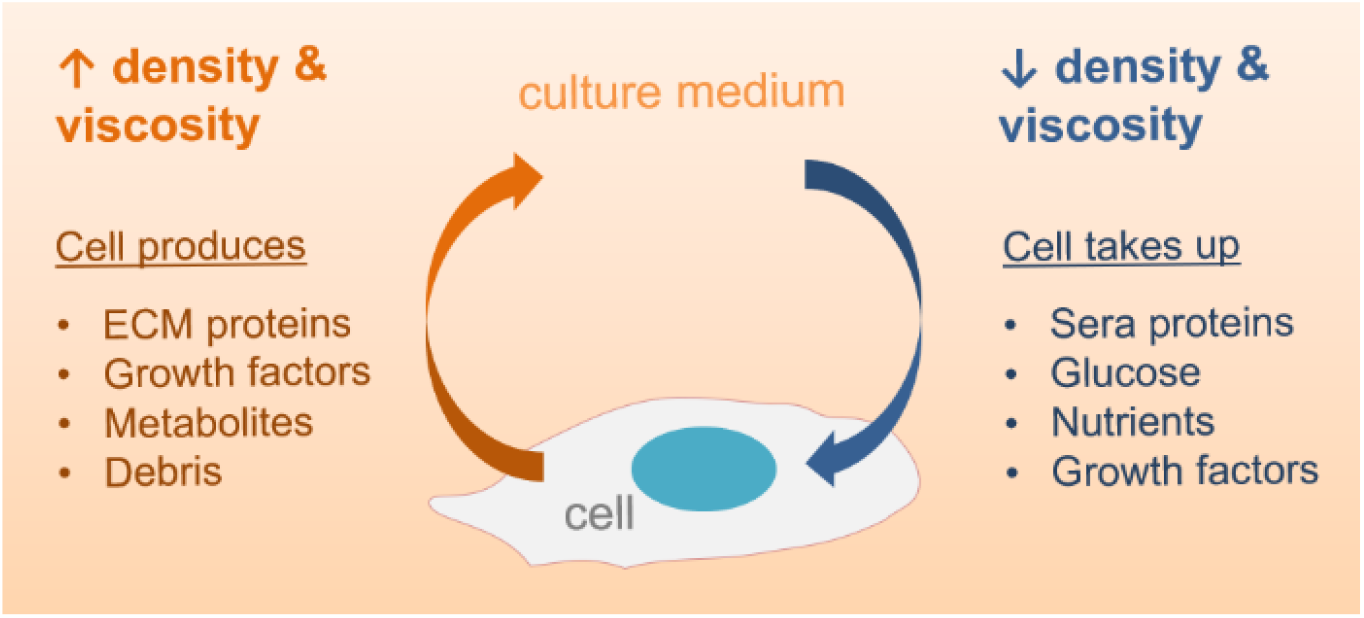
**The net fluid properties of culture medium are determined by the mass transport balance between cells and medium**

Given that the accuracy of any CFD simulation is directly determined by the assumptions, mathematical models, boundary conditions and fluid properties assigned during the setup of the model, it is important to use study-specific properties of culture media for more accurate CFD analysis of *in vitro* culture systems. It is understood that not all researchers have ready access to a rheometer or viscometer. Therefore, the densities and dynamic viscosities of two widely-used commercial culture media – Dulbecco’s Modified Eagle Medium (DMEM) and RPMI-1640 supplemented with 0, 5, 10 and 20% (v/v) foetal bovine serum (FBS) - were measured to serve as a reference for sera-free media (0%) and media supplemented with concentrations of FBS typically used for cell culture. This study also sought to investigate whether the physical properties of culture medium change during routine cell culture and found that both the density and viscosity of the medium increased over three days of standard passaging of HN6 human head and neck and NCI-H460 human lung carcinoma cell lines, with distinct differences between the results. Finally, CFD analysis of a simple microfluidic channel was conducted to investigate and compare any differences between modelling culture medium as water versus using experimentally-measured properties of culture medium.

## 2. Materials & Methods

### 2.1 Sample preparation & cell culture

10 mL samples of RPMI-1640 (Gibco, Thermo Fisher Scientific) and DMEM high glucose medium (11965-092, Gibco, Thermo Fisher Scientific) supplemented with 0, 5, 10 and 20% v/v FBS (10091148, Gibco, Thermo Fisher Scientific) were dispensed into capped tubes. To evaluate any changes in the density and viscosity of culture medium during cell culture, 10 mL samples were extracted during routine passaging of NCI-H460 and HN6 cells after 3 days of culture. Both cell lines were passaged with an initial seeding density of approximately 2 × 10^6^ cells per flask (T75) and grown in 12 mL of RPMI-1640 medium supplemented with 5% FBS in a standard incubator (37°C and 5% CO2) to approximately 70-80% confluency (approximately 4.5 × 10^6^ cells/mL for both cell lines) at day 3. All media samples were kept refrigerated then warmed up to 37°C in a water bath prior to measurement. Deionized water (ddH2O) and 1X phosphate buffer solution (PBS) controls were also prepared.

### 2.2 Density measurement

1 mL samples were initially dispended in a dish using a standard 1000 µL pipette and weighed on a digital laboratory scale (Sartorius BP210D, Goettingen, Germany). However, small droplets of culture media were found to adhere to the inner surface of the pipette tips which compromised the accuracy of the results. Therefore, subtractive measurements were taken. Briefly, individual sample tubes were placed on the scale, which was then tared. The tube was removed and vortexed for 5 seconds to ensure even solute distribution, then 1 mL was pipetted from the tube and conserved in another tube for rheological measurement. The original sample tube was then re-weighed and the resultant weight difference was recorded. Ten measurements were taken for each sample, using the same pipette for consistency. The fluid density for each sample was then calculated by dividing the weight values by 1 cm^3^, then averaged, collated and plotted for comparison. One-way analysis of variance (ANOVA) and a paired t-test were performed to determine any statistical significance between results.

### 2.3 Rheology

Fresh FBS-supplemented DMEM and RPMI-1640 culture media and spent media samples were prepared as in Section 2.1 along with deionized water controls. A rheometer (Paar Physica MCR 300 Modular Compact Rheometer, Anton Paar GmbH, Austria) was set up with parallel 50 mm diameter stainless steel plates at a horizontal position (0°C) and z-gap of 0.5 mm to define a sample volume of 0.981 mL. An adjunctive temperature control chamber was set to 37°C. 1 mL samples were individually loaded onto the centre of the lower platen, visible bubbles were removed with a syringe needle, then the top plate was lowered onto the sample. Sample fluid exceeding the test volume was trimmed with tissue paper. Rotational shear was then applied from 0 – 100 s^-1^. Measurements were conducted at 37°C with a minimum of 6 replicates per sample. The plates were thoroughly cleaned with 70% ethanol solution between each measurement and deionized water controls were used to calibrate the system between sample changes. Data was recorded on RheoCompass then exported for analysis. The datasets for each sample were plotted as viscosity profiles and shear stress-shear rate plots. The dynamic viscosity of each sample was then calculated by extracting the gradients of the shear-stress-shear plots.

### 2.4 Computational Fluid Dynamics

CFD simulations were conducted to determine any differences in flow mechanics produced within a simple microfluidics channel using experimentally-derived fluid properties for culture media compared to water. For this study, the values measured for fresh RPMI-1640 medium supplemented with 0, 5 and 10% v/v FBS and spent medium (originally RPMI-1640 supplemented with 5% FBS) after 3 days of culturing NCI-H460 and HN6 cells were modelled. These concentrations were selected to represent sera-free media and the most commonly used concentrations of FBS, as well as to determine and compare any differences in flow properties during/after cell culture. Briefly, a simple bifurcated microfluidic channel model (channel width = 1 mm, height = 0.5 mm, overall length = 20 mm) was created on SolidWorks (version 2019) then imported into ANSYS Fluent (version 2020 R2) for analysis. A tetrahedral dropped node mesh was generated from the channel geometry, which represents the fluid domain, and mesh refinement was performed to ensure adequate discretization. A mesh quality of 0.33 was confirmed in ICEM. The following assumptions were made and applied: (i) culture medium is an incompressible Newtonian fluid, (ii) steady state fully laminar flow (Reynolds number < 2000), (iii) flow occurs at 37°C at atmospheric pressure and a gravitational acceleration of 9.81m/s^2^ and (iv) non-slip wall boundaries. A steady state viscous laminar model with gravitational acceleration of 9.81m/s^2^ and a pressure-based solver at gauge pressure were selected. The channel inlet was set as a velocity inlet with directional flow defined via Cartesian co-ordinates in the negative Y direction (downwards into the fluid domain). A flow rate of 1.2 × 10^−6^ m/s was then applied to model the average physiological capillary perfusion rate [39] and the outlet was set at gauge pressure without backflow. The convergence criterion was set at 10^−7^ for continuity, x, y and z-velocities. To model different culture media, custom entries were created in the materials library using the density and viscosity values measured (Sections 3.1, 3.2). For this study, 5% and 10% v/v FBS-supplemented RPMI-1640 media and spent media after culturing cells (originally 5% FBS-supplemented RPMI-1640) were modelled, along with default water properties and water at 37°C. The cell zone was set to ‘fluid’ and the experimentally-derived culture medium properties were assigned. After model setup, the respective material properties of each fluid (Table 1) were applied for simulation.

**Table 1.**
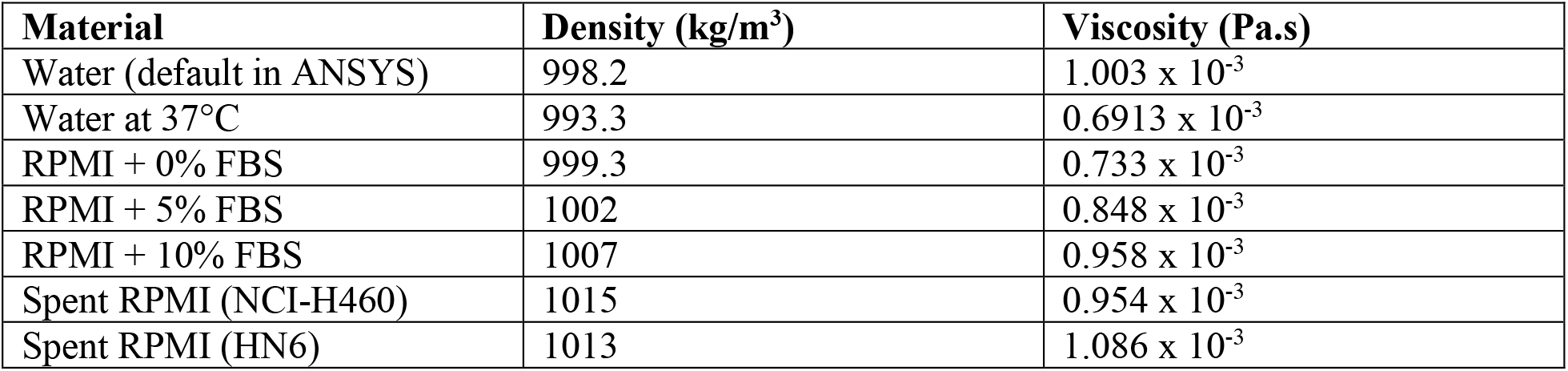
Properties of water and different culture media formulations assigned to CFD model of a microfluidics channel.

| Material | Density (kg/m<sup>3</sup>) | Viscosity (Pa.s) |
| --- | --- | --- |
| Water (default in ANSYS) | 998.2 | 1.003 x 10<sup>-3</sup> |
| Water at 37°C | 993.3 | 0.6913 x 10<sup>-3</sup> |
| RPMI + 0% FBS | 999.3 | 0.733 x 10<sup>-3</sup> |
| RPMI + 5% FBS | 1002 | 0.848 x 10<sup>-3</sup> |
| RPMI + 10% FBS | 1007 | 0.958 x 10<sup>-3</sup> |
| Spent RPMI (NCI-H460) | 1015 | 0.954 x 10<sup>-3</sup> |
| Spent RPMI (HN6) | 1013 | 1.086 x 10<sup>-3</sup> |

Simulations were then performed with a maximum of 1000 iterations or until convergence. Plots of wall shear stress, global pressure, velocity vectors and flow path lines were generated in CFD-Post to visualize flow field properties.

## 3. Results & Discussion

### 3.1 Gravimetric density of culture media

The density of a material is a defined as its mass per unit volume by the formula ρ = mass/volume, with units expressed in g/cm^3^ or kg/m^3^ (1 mL = 1 cm^3^ for fluids). Density is influenced by temperature and pressure, which affect the volume and molecular state of a material. As the intended operational conditions for the majority of *in vitro* culture devices occur at atmospheric pressure (1 atm) and 37°C, the culture media samples and controls were measured under these conditions.

#### 3.1.1 Methodology

The subtractive measurement approach was straightforward and addressed the issue of uncontrolled droplet retention within the pipette tip that was initially encountered when dispensed volumes of culture medium were weighed.. The charge density of the proteins, salts and sugars present in the media samples likely contributed to higher surface tension which led to sample retention within the pipette tips. Otherwise, no droplets were observed on the outer surface of the tips when drawing samples during subtractive weighing. This method is therefore recommended for measuring the density of culture media and other ionically-charged and/or colloidal solutions.

#### 3.1.2 Results

The densities of the culture media samples and controls are presented in Figure 2 and Table 2 as follows:

**Table 2.** Density and dynamic viscosity values measured for each sample.

| %FBS (v/v) | Density (g/cm<sup>3</sup>) |  | Dynamic Viscosity (mPa.s) |  |
| --- | --- | --- | --- | --- |
|  | DMEM (high glucose) | RPMI-1640 | DMEM (high glucose) | RPMI-1640 |
| **0** | 1.000 ± 0.001 | 0.999 ± 0.003 | 0.731 ± 0.015 | 0.733 ± 0.006 |
| **5** | 1.002 ± 0.003 | 1.002 ± 0.002 | 0.862 ± 0.029 | 0.848 ± 0.021 |
| **10** | 1.009 ± 0.003 | 1.007 ± 0.002 | 0.930 ± 0.034 | 0.958 ± 0.032 |
| **20** | 1.023 ± 0.005 | 1.020 ± 0.005 | 1.050 ± 0.027 | 1.089 ± 0.044 |
| **Water** | | 0.995 ± 0.002 | 0.659 ± 0.017 | |
| **PBS** | | 0.998 ± 0.002 | N/A | |
| **NCI-H460 spent medium** | | 1.015 ± 0.003 | 0.954 ± 0.070 | |
| **HN6 spent medium** | | 1.013 ± 0.003 | 1.086 ± 0.073 | |

**Figure 2.**
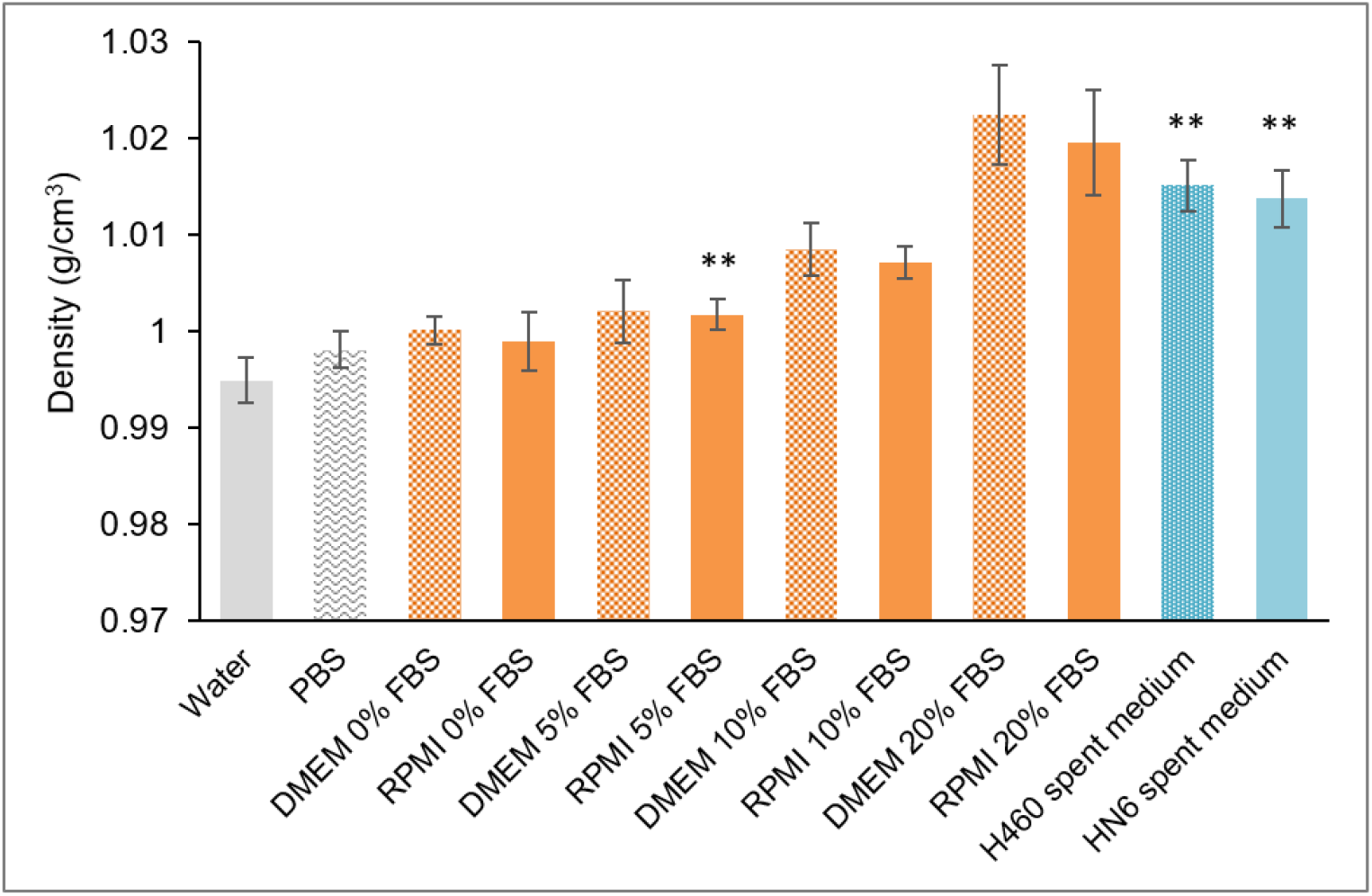
**The density of DMEM (high glucose) and RPMI-1640 media supplemented with 0, 5, 10, 20 % v/v FBS and 5 % FBS-supplemented RPMI-1640 media after 3 days of standard subculture of NCI-H460 and HN6 cell lines. Deionized water and PBS were measured as controls. Values are expressed as mean ± standard deviation. Significance (p < 0.01) from a paired t-test and one-way ANOVA as shown.**

#### 3.1.3 Discussion

The density of water at 37°C is 0.9933 g/cm^3^ (IAPWS R12-08), hence the value measured for the deionized water control (0.995 g/cm^3^) is within acceptable calibration standards for the 1000 μL pipette (Gibson). All culture media samples and 1X PBS were found to have higher densities than deionized water (Figure 2), which was as expected due to the presence of salts, sugars and other solutes within these solutions. Significant differences (p<0.01) were found between the densities measured for all culture media solutions, PBS and water. The density of both DMEM and RPMI-1640 medium increased directly proportional to the volume of added FBS (Figure 2), which was as expected given that FBS is the largest % volume additive in complete media and the main component of FBS is bovine serum albumin, a heavy, high molecular weight protein. High glucose DMEM, which contains 4.5 g/L of glucose, was consistently found to have higher density than RPMI-1640, which contains 2.0 g/L glucose. Otherwise, both media formulations contain approximately 11 g of inorganic salts and the relatively smaller percentage volumes of other additives such as growth factors can be assumed to have negligible effect on fluid properties.

The net density of 5% FBS-supplemented RPMI-1640 media was found to have increased significantly (p < 0.01) and by 1.297% and 1.197% after culturing NCI-H460 and HN6 cell lines for 3 days respectively, supporting the axiomatic hypothesis that general cellular activities including metabolism, secretion of ECM proteins, factors, signalling molecules, waste products and debris increase the net density of culture medium during culture. Differences between the results for the NCI-H460 and HN6 cell lines can be attributed to cell type-specific metabolic and proliferation rates, as well as the quantity, type, size and other properties of the proteins secreted by each respective cell line. This increase in fluid density equates to higher hydrostatic pressure P as given by P = ρgh, where ρ is the density, g denotes the gravitational constant and h is the depth of the fluid. Any changes in pericellular pressure may be significant or is at least perceived on a cellular level, but it is likely that changes in osmotic flux and oxygen diffusion in the medium due to increased ionic concentration from cellular secretion would have a greater effect on cellular responses. The density values were then applied in preliminary CFD analysis of a simple microfluidic organ-on-chip channel.

### 3.2 Rheological properties of culture medium

Rheometry was performed to measure the dynamic viscosity of the culture media samples. Viscosity corresponds to the informal concept of ‘thickness’ and provides a measure of the resistance of a fluid to deformation or flow with application of shear or tensile stress, as defined by the following equation:

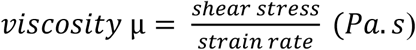

Viscosity is a quantity expressing the magnitude of internal friction within a fluid, i.e. intermolecular friction between neighbouring particles that are moving at different velocities. The size, shape and magnitude of intermolecular forces of particles within a fluid determine its viscosity. As cell cultures proliferate, metabolic activity and other processes can be expected to increase the number of solute particles in culture media, thereby increasing its viscosity. For most liquids, viscosity decreases as temperature increases and rises with pressure as inter-particle energies are temperature and pressure-dependent. Therefore, samples were analyzed at 37°C and 1 atm for model fidelity as for density measurement.

#### 3.2.1 Methodology

The protocol described is standard for conducting rheometry with a parallel plate configuration, as appropriate for measuring low to medium viscosity fluids. A faint residue was observed on the plates after each sample measurement, indicating potential adsorption of proteins and molecules onto the test apparatus which may affect the consistency of the results. Although care was taken, other potential sources of error include the presence of microbubbles introduced during sample loading, inconsistencies in sample volume in the order of µL from sample trimming, test temperature stability and accidental contamination of the samples with particulate matter such as dust.

#### 3.2.2 Results

The dynamic viscosities of each sample are presented in Figure 3 as follows.

**Figure 3.**
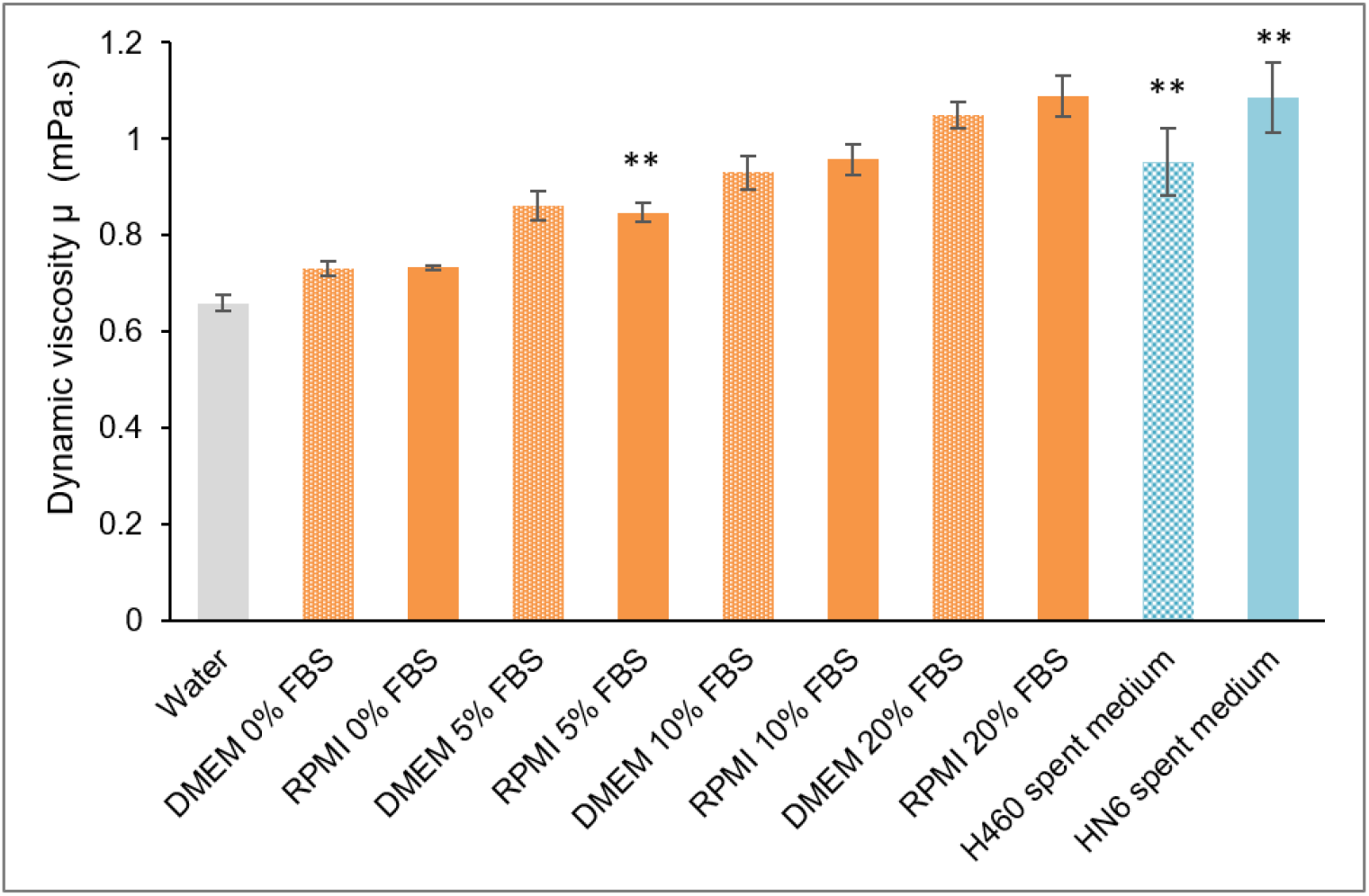
**Dynamic viscosities of DMEM (high glucose) and RPMI-1640 media supplemented with typical concentrations of FBS (0, 5, 10, 20% v/v) and 5% FBS-supplemented RPMI-1640 after 3 days of standard subculture of NCI-H460 and HN6 cell lines. Samples were measured at 37°C along with a deionized water control. Values are expressed as mean ± standard deviation. Significance (p value < 0.01) from one-way ANOVA and paired t-test as shown.**

#### 3.2.3 Discussion

Culture media are frequently modelled as Newtonian fluids in CFD analyses of *in vitro* culture systems [26, 36, 40-42]. Newtonian fluids have a constant viscosity that is independent of shear stress and strain rate, which is exhibited as a linear trend on shear stress-shear rate plots and viscosity-shear profiles. Linear trends (R^2^ values > 0.995) were obtained for all media samples including the deionized water controls (Supplementary Data 1), indicating that DMEM and RPMI-1640 media supplemented with typical concentrations of FBS can be considered as Newtonian fluids for simplified computational modelling purposes. However, slight shear thinning was observed at lower shear rates (0-20 s^-1^) on all viscosity-shear rate profiles, which became more pronounced with higher concentrations of added FBS (Supplementary data 2). This is consistent with the shear thinning previously reported for culture media with added sera proteins [43, 44], albumin [44, 45] and protein-rich biological fluids such as blood plasma [46-48]. Higher concentrations of FBS than those investigated in this study or addition of viscosity modifiers such as dextran may produce more distinctively non-Newtonian fluid behaviours due to the effects of shear-induced polymer chain alignment [49]. It must be noted that some measurement instability was observed on the viscosity profiles at lower shear rates up to 20 s^-1^, particularly for water and media samples with lower concentrations of added FBS (Supplementary Material 2). This can be attributed to the inherent sensitivity of the rheometer and test apparatus. While the viscosities calculated from the shear stress-shear rate profiles are valid, any shear thinning (non-Newtonian fluid) behaviour warrants particular consideration for devices designed to apply low flow rates to cell cultures. It is recommended that measurements be taken with more sensitive instrumentation to further elucidate the rheology of culture media at lower shear rates as otherwise shear stresses may be significantly underestimated in CFD simulation, which in turn can inaccurately inform the selection and application of suboptimal flow rates as well as obfuscate correlations between shear estimates and cellular responses

The average dynamic viscosity of the deionized water control was measured to be 0.659 mPa.s (Figure 3, Table 2), which was lower than the 0.6913 mPa.s standard for water at 37°C (IAPWS R12-08). The difference is not statistically significant and may potentially be due to thermal overshooting within the temperature control chamber i.e. the test temperature may have been closer to 38°C instead of 37°C and hence the viscosity was lower. Nevertheless, distinct viscosities were measured for each sample, where.all culture media samples were confirmed to be more viscous than the deionized water control. As with density, the dynamic viscosity of the media increased proportional to the % volume of added FBS(Figure 2, Figure 3). Even a 5% (v/v) addition of FBS, the lowest concentration, yielded a 30% increase in viscosity for both culture media compared to water at the same temperature, while the viscosities of 10% and 20% FBS-supplemented RPMI-1640 media was found to be 48% and 107% higher than that of the water control respectively (Table 3). Importantly, 5% FBS-supplemented RPMI media was found to have increased by 12.08% and 39.22% after 3 days of culture of NCI-H460 and NH6 cell lines respectively (Table 3). This change can be significant on a cellular level, where viscosity impacts molecular diffusion kinetics (passive mass transport) and a higher fluid viscosity equates to higher hydrostatic forces and shear stresses exerted on cells in flow culture devices such as organ-on-chips, which affect cell motility, cell-substrate interactions and migration [50].

**Table 3.** Dynamic viscosities of culture media and % increase compared to water.

| Sample (at 37°C) | Dynamic viscosity $\mu$ (mPa.s) | % increase compared to water |
| --- | --- | --- |
| Water | 0.659 ± 0.017 | 0 |
| DMEM + 0% FBS | 0.731 ± 0.015 | 10.924 |
| DMEM + 5% FBS | 0.861 ± 0.029 | 30.706 |
| DMEM + 10% FBS | 0.930 ± 0.034 | 41.069 |
| DMEM + 20% FBS | 1.050 ± 0.027 | 59.250 |
| RPMI-1640 + 0% FBS | 0.733 ± 0.006 | 11.138 |
| RPMI-1640 + 5% FBS | 0.848 ± 0.021 | 28.597 |
| RPMI-1640 + 10% FBS | 0.958 ± 0.032 | 45.255 |
| RPMI-1640 + 20% FBS | 1.089 ± 0.044 | 65.135 |
| NCI-H460 (RPMI-1640 + 5% FBS) | 0.954 ± 0.070 | 44.619 |
| HN6 (RPMI-1640 + 5% FBS) | 1.086 ± 0.074 | 64.656 |

The magnitude and rate of change in the fluid properties of culture medium is affected by numerous experimental factors, including initial seeding density, duration of culture, the substrate, the media formulation itself (e.g. addition of growth factors), the geometry of the culture domain, flow conditions (e.g. pulsatile), percentage and rate of recirculation as well as the location of cells within the system. In addition, there is an interdependency where the density and viscosity of culture media directly affects the diffusivity of soluble molecules and determine the magnitude of hydrostatic pressure (mechanical stimuli) exerted by the fluid environment on cells, which affect their function. Cell-typespecific characteristics will also determine the viscosity of the culture medium, including the types and quantities types of proteins secreted and mechanosensitivity to the effects of shear in flow-culture devices. Together, these factors collectively determine the diffusion profile of metabolites and secreted ECM proteins conceptually illustrated in Figure 1.

In devices where fresh medium is continually perfused, diffusion kinetics becomes a function of the flow rate, whereas in systems that recirculate medium, recirculation introduces temporal effects where as cells grow, the balance between continual nutrient depletion and increasing concentrations of metabolites both affect and occur as a function of the proliferation rate. In perfused devices, fluid flow introduces mixing effects that reduce stagnant regions where heavier proteins and solutes can sink or aggregate, and so theoretically produces a more homogeneous fluid density throughout the system. Furthermore, introduction of perfusion provides mechanical stimuli and improves mass transport, factors which are known to enhance cell growth and proliferation [34, 51]. More cells equate to higher quantities of secreted factors and proteins and consequently the physical properties of the culture medium would increase over time in closed/recirculated systems. The magnitude and rate of increase must be experimentally verified for each system. Cell culture suspensions such as those cultured in bioreactors would have different rheological properties that likewise require characterization due to differences in cell size, shape, deformability, concentration and size of aggregates [52]. Finally, although the overall range of viscosity values and rheological trends measured here can be expected to be similar for commercially available culture media, supplier and batch-to-batch variations and experimental modifications e.g. conditioning steps may affect the consistency of resultsof, thus further recommending study-specific characterization.

Overall, these results demonstrate that the dynamic viscosities of DMEM and RPMI-1640 media supplemented with conventional concentrations of FBS are significantly higher than that of water at 37°C. Given that viscosity is a key property that determines fluid behaviour, these results recommend using experimentally-derived fluid properties for CFD analysis where possible. Differences in hydrodynamics generated by these experimentally derived culture medium properties were then compared to water models by CFD analysis.

### 3.3 CFD models using experimentally-derived fluid properties

CFD simulations were performed to examine and compare any differences between models that apply the default properties of water typically provided in ANSYS versus using actual values for density and viscosity of water at 37°C and culture media that were measured experimentally. Results and implications for cell cultures are presented and discussed as follows.

#### 3.3.1 Results

The wall shear stress and global pressure profiles across the channel model and maximum values of these variables are presented and summarised in Table 4 and Table 5 as follows:

**Table 4.**
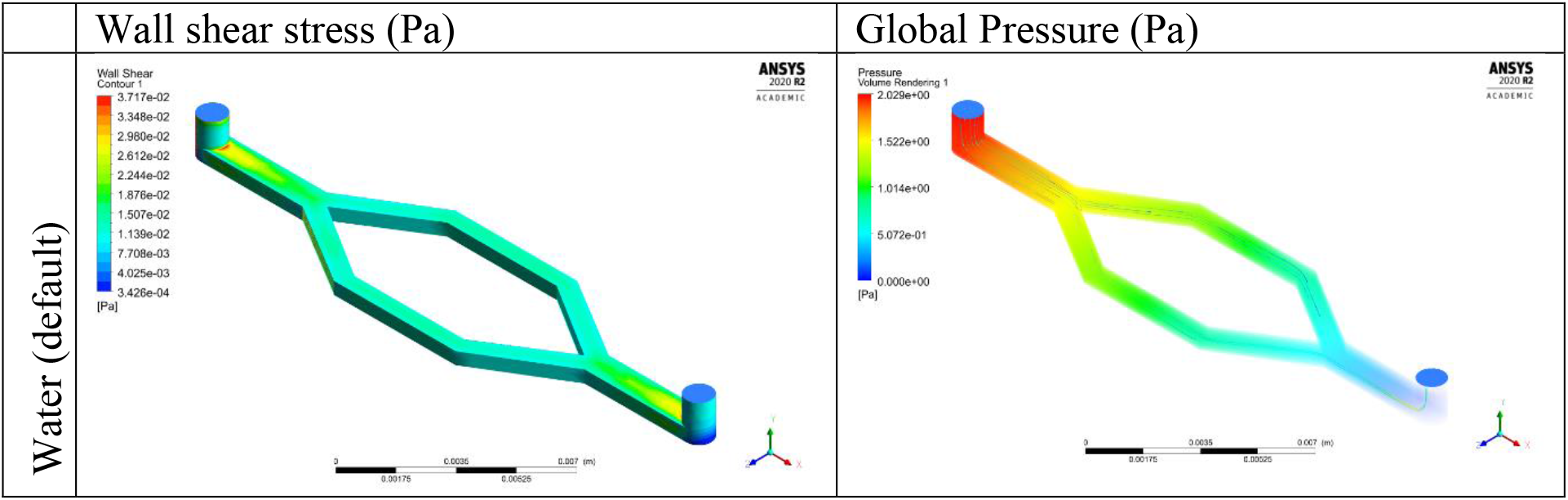
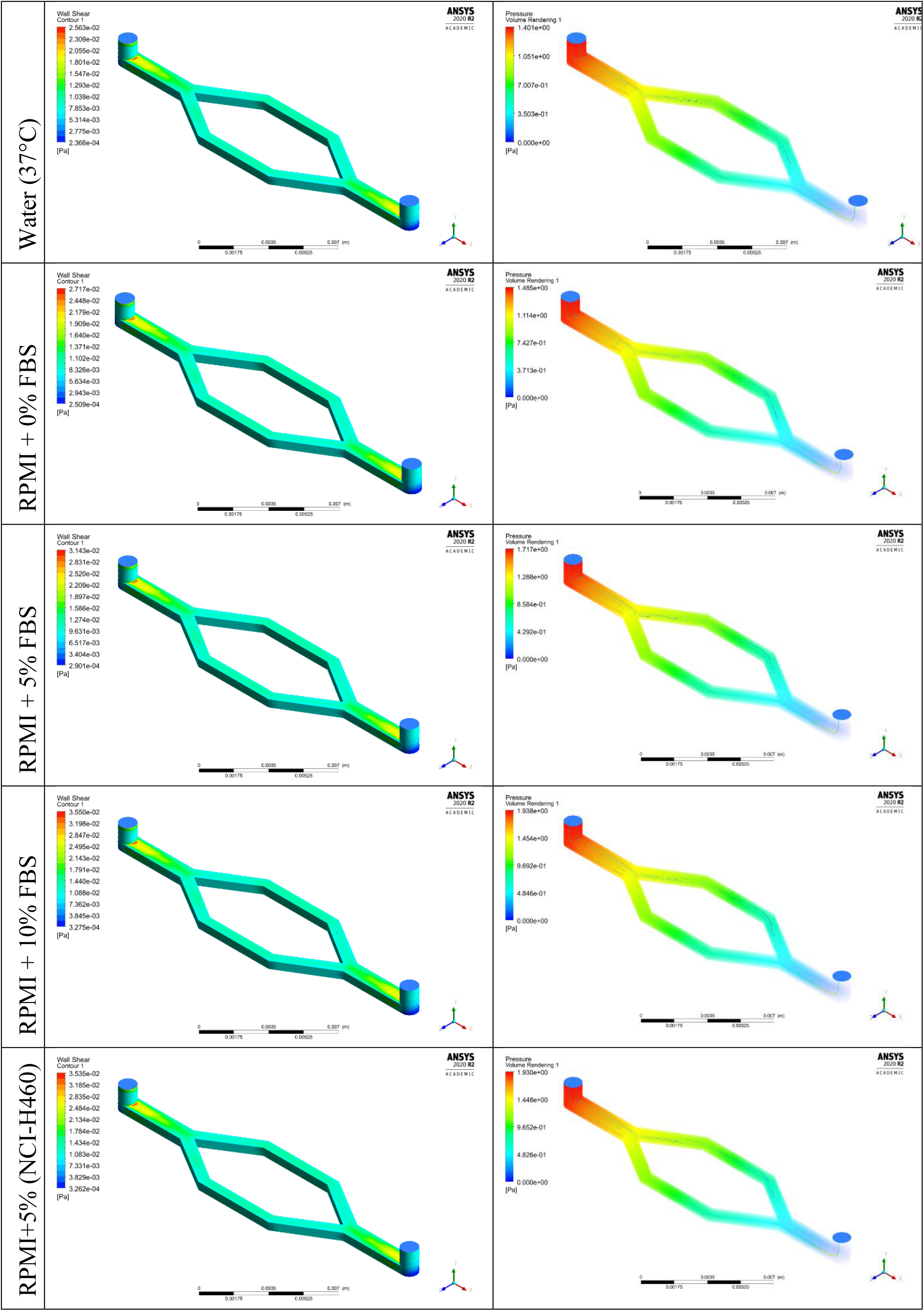
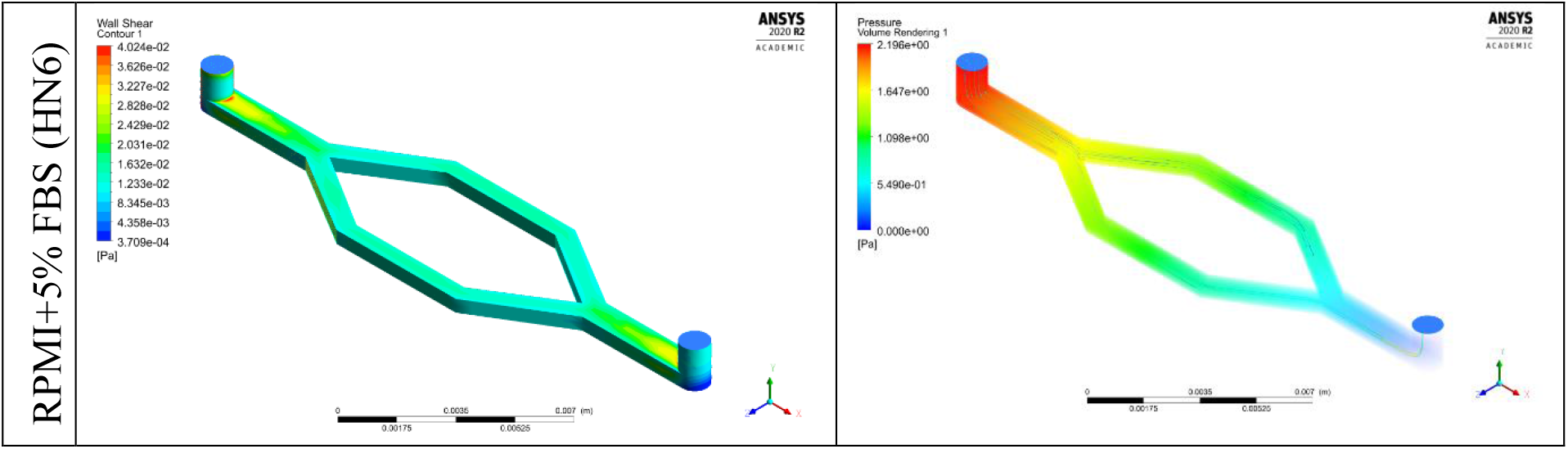
Wall shear stress and global pressure distributions within the channel model.

**Table 5.** Maximum wall shear stress and pressure for each model.

| Model | Max Wall Shear Stress (mPa) | Max Pressure (Pa) |
| --- | --- | --- |
| Water (default) | 37.17 | 2.029 |
| Water (37°C) | 25.63 | 1.401 |
| RPMI-1640 + 0% FBS | 27.17 | 1.485 |
| RPMI-1640 + 5% FBS | 31.43 | 1.717 |
| RPMI-1640 + 10% FBS | 35.50 | 1.938 |
| Spent RPMI-1640* (H460) | 35.35 | 1.930 |
| Spent RPMI-1640* (HN6) | 40.24 | 2.196 |

\*supplemented with 5% FBS

#### 3.3.2 Discussion

The purpose of this study was to investigate differences between CFD models using experimentally derived properties of culture media and water. Hence although rheological results indicated that culture media is slightly shear thinning at lower shear rates (Section 3.2, Supplementary material), culture media were modelled as Newtonian fluids for simplicity and comparability with existing studies that apply Newtonian fluid assumptions. The corresponding shear rate for maximum wall shear stress is approximately 37 s^-1^ for all models (Table 5). However, the main cell culture region (within the bifurcated channels for this particular model) would experience shear rates approximately between 10-20 s^-1^ for the flow rate investigated (Table 4), which is within the range where shear thinning behaviour was observed on viscosity profiles (Supplementary Material 2). Therefore, it is possible that shear stresses may be underestimated should viscosities be higher at lower shear rates, and so cells may be exposed to damaging levels of shear should flow rates be informed from simulation. Applying Weissenberg-Rabinowitsch corrections and Sisko fluid model assumptions [43] may be necessary to achieve more accurate simulations, particularly at lower flow rates.

CFD simulations provide an estimate of the upper bounds of the range of shear stresses within a fluid system as well as the locations of maximum stress and pressure, which are important to know and identify in order to assess the risk of failure due to rupture and crucially for *in vitro devices*, whether and where cells may encounter shear damage at a particular flow rate. As can be seen in the shear stress profiles (Table 4), the location of the maximum wall shear stress occurred at the inner junction between the inlet and outlet ports (cylindrical features) and the channel for all models at the flow rate investigated; this was as expected given that the flow rate is relatively low and the fluid properties measured are sufficiently similar (Table 2). Overall, the shear stress values obtained from simulation can be considered to be moderate i.e. unlikely to cause shear damage to cells [53], and are within a range previously reported to be suitable for promoting myogenesis [54], glial proliferation [55] and osteogenic differentiation [56, 57]. Global pressure profiles were similarly consistent across all models given due to the flow conditions originally assigned. Wall shear stress and global pressure increased proportional to the dynamic viscosity and density of the fluid in accordance with the Navier Stokes equations (Equation 1), where higher % volumes of FBS (higher viscosity) directly corresponded with higher shear and pressure within the channel (Table 2, Table 5). The maximum wall shear stress of 5% FBS supplemented RPMI-medium increased by 12.5% and 28.0% after 3 days of culturing NCI-H460 and HN6 cell lines respectively (Table 5). This result highlights the importance of determining model-specific fluid properties, particularly for analyzing systems where culture medium is recirculated.

Interestingly, the default properties of water in ANSYS yielded results similar to those of RPMI-1640 medium supplemented with 10% FBS and the spent medium (Table 5) as the viscosity was closer to values measured for media with at least 10% v/v added FBS and spent media (Table 1, Table 2). While this indicates that computational studies that have applied default water properties (typically at room temperature) in simulation programs can still be considered to be valid, it is possible that actual viscosities of culture media are higher at lower shear rates and so corresponding shear experienced by cells may be up to orders of magnitude higher than values estimated through CFD simulation. Nevertheless, the accuracy of any CFD model depends on inputs andit is ideal to use actual experimentally-derived properties of culture medium, such as those measured in this work. For further model accuracy, measuring fluid properties over a time course and the surface roughness of substrate materials e.g. polycarbonate, glass, PDMS in their in situ experimental state (i.e. including any coatings or treatments) are recommended.

## Conclusions

This study sought to measure the dynamic viscosity and density of common media formulations to serve as a ballpark reference for CFD analysis of cell and tissue culture devices involving flow. In this work, the density and dynamic viscosity of high glucose DMEM and RPMI-1640 media supplemented with typical concentrations of FBS (0, 5, 10 and 20% v/v) were measured. RPMI-1640 media after 3 days of culture of two cell lines was also measured as a preliminary investigation of how the physical properties of culture medium change during cell culture. The densities and dynamic viscosities of all media samples were definitively shown to be higher than that of PBS and deonized water at 37°C, where density and viscosity increased directly proportional to concentration of added FBS. Importantly, both the density and viscosity of 5% FBS-supplemented RPMI-1640 were found to increase significantly after 3 days of culture of NCI-H460 and HN6 cell lines, where distinct differences between the results for each cell line indicate cell line and model specificity. As previously discussed, the magnitude and rate of change in these properties varies between cell types and numerous other factors including initial seeding density. For this reason, conducting a time course to determine time-resolved changes in the viscosity and density of culture medium during the course of culture was beyond the scope of this study. However, it is recommended and will be performed for validation studies.

CFD simulations of physiological flow through a simple organ-on-chip channel using the density and viscosity values measured demonstrated differences in the magnitudes of wall shear stress and global pressure between different media formulations (0, 5, 10% added FBS) and a water model, where wall shear stress and pressure increased with higher concentrations of added FBS (higher viscosity). Both maxima and global wall shear stress and pressure significantly increased when modelled with properties of medium after3 days of culture of NCI-H460 and HN6 cell lines, bearing implications for systems that recirculate medium over any period of culture such as bioreactors Applying the default values for water in ANSYS yielded results similar to those of RPMI-1640 medium supplemented with at least 10% FBS. Therefore, studies that have modelled culture medium as water and applied density and dynamic values within the ranges measured can still be considered as valid. Nevertheless, these results highlight the importance of using model-specific fluid properties for CFD analysis and recommend the use of experimentally-derived properties of culture media for computational analysis of cell and tissue culture devices. It is hoped that the values measured for DMEM and RPMI-1640 media will serve as a useful baseline reference for anyone conducting CFD analyses of cell or tissue culture devices. It must be noted that these values are most applicable for systems where cells are attached to a substrate i.e. not suspended in the medium. Cell suspensions would have different rheological properties and require characterization. Time-resolved measurements are likewise necessary for systems where medium is recirculated for any duration of culture. Overall, this work recommends determining both baseline and temporal fluid properties of culture media for more accurate CFD analysis and optimized design of *in vitro* culture systems.

## Acknowledgements

The author would like to thank the Melanoma Institute Australia and the Department of Infectious Diseases (The University of Sydney) for providing the culture media samples tested in this study.

## References

[1] T. Yao and Y. Asayama, “Animal-cell culture media: History, characteristics, and current issues,” (in eng), Reproductive medicine and biology, vol. 16, no. 2, pp. 99–117, 2017.

[2] A. Sen, M. S. Kallos, and L. A. Behie, “Expansion of mammalian neural stem cells in bioreactors: effect of power input and medium viscosity,” Developmental Brain Research, vol. 134, no. 1–2, pp. 103–113, 2002.

[3] J. L. Moreira et al., “Effect of Viscosity upon Hydrodynamically Controlled Natural Aggregates of Animal Cells Grown in Stirred Vessels,” Biotechnology Progress, vol. 11, no. 5, pp. 575–583, 1995.

[4] S. H. Cartmell, B. D. Porter, A. J. García, and R. E. Guldberg, “Effects of Medium Perfusion Rate on Cell-Seeded Three-Dimensional Bone Constructs in Vitro,” Tissue Engineering, vol. 9, no. 6, pp. 1197–1203, December 2003 2003.

[5] R. Pörtner and C. Giese, “An Overview on Bioreactor Design, Prototyping and Process Control for Reproducible Three-Dimensional Tissue Culture,” no. 3, pp. 53–78, 2007.

[6] J. Rouwkema, S. Gibbs, M. P. Lutolf, I. Martin, G. Vunjak-Novakovic, and J. Malda, “In vitro platforms for tissue engineering: implications for basic research and clinical translation,” Journal of tissue engineering and regenerative medicine, vol. 5, no. 8, pp. e164–e167, 2011.

[7] T. Gareau et al., “Shear stress influences the pluripotency of murine embryonic stem cells in stirred suspension bioreactors,” (in eng), J Tissue Eng Regen Med, vol. 8, no. 4, pp. 268–78, Apr 2014.

[8] W. J. Polacheck, R. Li, S. G. M. Uzel, and R. D. Kamm, “Microfluidic platforms for mechanobiology,” (in eng), Lab on a chip, vol. 13, no. 12, pp. 2252–2267, 2013.

[9] H. W. Hou, W. C. Lee, M. Leong, S. Sonam, S. Vedula, and C. T. Lim, “Microfluidics for Applications in Cell Mechanics and Mechanobiology,” Cellular and Molecular Bioengineering, vol. 4, pp. 591-602, 01/01 2012.

[10] F. Kurth, K. Eyer, A. Franco-Obregón, and P. S. Dittrich, “A new mechanobiological era: microfluidic pathways to apply and sense forces at the cellular level,” Current Opinion in Chemical Biology, vol. 16, no. 3, pp. 400-408, 2012/08/01/ 2012.

[11] E. Ergir, B. Bachmann, H. Redl, G. Forte, and P. Ertl, “Small Force, Big Impact: Next Generation Organ-on-a-Chip Systems Incorporating Biomechanical Cues,” (in English), Frontiers in Physiology, Mini Review vol. 9, no. 1417, 2018-October-09 2018.

[12] J. M. Osborne, R. D. O’Dea, J. P. Whiteley, H. M. Byrne, and S. L. Waters, “The Influence of Bioreactor Geometry and the Mechanical Environment on Engineered Tissues,” Journal of Biomechanical Engineering, vol. 132, no. 5, 2010.

[13] S. G. Mina, W. Wang, Q. Cao, P. Huang, B. T. Murray, and G. J. Mahler, “Shear stress magnitude and transforming growth factor-βeta 1 regulate endothelial to mesenchymal transformation in a three-dimensional culture microfluidic device,” RSC Advances, 10.1039/C6RA16607E vol. 6, no. 88, pp. 85457–85467, 2016.

[14] J. M. Barnes, J. T. Nauseef, and M. D. Henry, “Resistance to Fluid Shear Stress Is a Conserved Biophysical Property of Malignant Cells,” PLOS ONE, vol. 7, no. 12, p. e50973, 2012.

[15] G. Kretzmer and K. Schügerl, “Response of mammalian cells to shear stress,” Applied Microbiology and Biotechnology, vol. 34, no. 5, pp. 613-616, 1991/02/01 1991.

[16] W. Yu et al., “A Microfluidic-Based Multi-Shear Device for Investigating the Effects of Low Fluid-Induced Stresses on Osteoblasts,” PLOS ONE, vol. 9, no. 2, p. e89966, 2014.

[17] F. Tovar-Lopez et al., “A Microfluidic System for Studying the Effects of Disturbed Flow on Endothelial Cells,” (in English), Frontiers in Bioengineering and Biotechnology, Brief Research Report vol. 7, no. 81, 2019-April-17 2019.

[18] J. G. Santiago, S. T. Wereley, C. D. Meinhart, D. J. Beebe, and R. J. Adrian, “A particle image velocimetry system for microfluidics,” Experiments in Fluids, vol. 25, no. 4, pp. 316-319, 1998/09/01 1998.

[19] A. Campos Marin, T. Grossi, E. Bianchi, G. Dubini, and D. Lacroix, “2D µ-Particle Image Velocimetry and Computational Fluid Dynamics Study Within a 3D Porous Scaffold,” Annals of Biomedical Engineering, vol. 45, no. 5, pp. 1341-1351, 2017/05/01 2017.

[20] H.-S. Chuang and Y.-L. Lo, “Microfluidic velocity measurement using a scanning laser Doppler microscope,” Optical Engineering, vol. 46, no. 2, p. 024301, 2007.

[21] Y. S. Morsi, W. W. Yang, A. Owida, and C. S. Wong, “Development of a novel pulsatile bioreactor for tissue culture,” (in eng), J Artif Organs, vol. 10, no. 2, pp. 109–14, 2007.

[22] L. Stern et al., “Doppler-based flow rate sensing in microfluidic channels,” (in eng), Sensors (Basel, Switzerland), vol. 14, no. 9, pp. 16799–16807, 2014.

[23] J. D. Salvi, J. Y. Lim, and H. J. Donahue, “Finite Element Analyses of Fluid Flow Conditions in Cell Culture,” Tissue Engineering. Part C, Methods, vol. 16, no. 4, pp. 661–670, 2010.

[24] D. Freitas, H. A. Almeida, and P. J. Bártolo, “Perfusion Bioreactor Fluid Flow Optimization,” Procedia Technology, vol. 16, pp. 1238-1247, 2014/01/01/ 2014.

[25] T. Glatzel et al., “Computational fluid dynamics (CFD) software tools for microfluidic applications – A case study,” Computers & Fluids, vol. 37, no. 3, pp. 218-235, 2008/03/01/ 2008.

[26] M. Huang, S. Fan, W. Xing, and C. Liu, “Microfluidic cell culture system studies and computational fluid dynamics,” Mathematical and Computer Modelling, vol. 52, no. 11, pp. 2036-2042, 2010/12/01/ 2010.

[27] A. Marturano-Kruik et al., “Human bone perivascular niche-on-a-chip for studying metastatic colonization,” Proceedings of the National Academy of Sciences, vol. 115, p. 201714282, 01/23 2018.

[28] A. R. Patrachari, J. T. Podichetty, and S. V. Madihally, “Application of computational fluid dynamics in tissue engineering,” Journal of bioscience and bioengineering, vol. 114, no. 2, pp. 123–32, 2012.

[29] B. S. Borys, E. L. Roberts, A. Le, and M. S. Kallos, “Scale-up of embryonic stem cell aggregate stirred suspension bioreactor culture enabled by computational fluid dynamics modeling,” Biochemical Engineering Journal, vol. 133, pp. 157-167, 2018/05/15/ 2018.

[30] D. W. Hutmacher and H. Singh, “Computational fluid dynamics for improved bioreactor design and 3D culture,” Trends in biotechnology, vol. 26, no. 4, pp. 166–72, 2008.

[31] M. Israelowitz, B. Weyand, S. Rizvi, P. Vogt, and H. von Schroeder, “Development of a Laminar Flow Bioreactor by Computational Fluid Dynamics,” Journal of Healthcare Engineering, vol. 3, pp. 455-476, 09/01 2012.

[32] M. Cioffi, F. Boschetti, M. T. Raimondi, and G. Dubini, “Modeling evaluation of the fluid-dynamic microenvironment in tissue-engineered constructs: a micro-CT based model,” Biotechnology and Bioengineering, vol. 93, no. 3, pp. 500–10, 2006.

[33] M. Malvè, D. J. Bergstrom, and X. B. Chen, “Modeling the flow and mass transport in a mechanically stimulated parametric porous scaffold under fluid-structure interaction approach,” International Communications in Heat and Mass Transfer, vol. 96, pp. 53-60, 2018/08/01/ 2018.

[34] S. Sugiura, Y. Sakai, K. Nakazawa, and T. Kanamori, “Superior oxygen and glucose supply in perfusion cell cultures compared to static cell cultures demonstrated by simulations using the finite element method,” Biomicrofluidics, vol. 5, no. 2, p. 022202, 2011.

[35] Y. Guyot, F. P. Luyten, J. Schrooten, I. Papantoniou, and L. Geris, “A Three-Dimensional Computational Fluid Dynamics Model Of Shear Stress Distribution During Neotissue Growth In A Perfusion Bioreactor,” Biotechnology and Bioengineering, vol. 112, 06/01 2015.

[36] B. Porter, R. Zauel, H. Stockman, R. Guldberg, and D. Fyhrie, “3-D computational modeling of media flow through scaffolds in a perfusion bioreactor,” Journal of Biomechanics, vol. 38, no. 3, pp. 543–9, 2005.

[37] S. P. Singh, M. Shukla, and R. K. Srivastava, “Lattice Modeling and CFD Simulation for Prediction of Permeability in Porous Scaffolds,” Materials Today: Proceedings, vol. 5, no. 9, Part 3, pp. 18879-18886, 2018/01/01/ 2018.

[38] S. Yedgar, D. B. Weinstein, W. Patsch, G. Schonfeld, F. E. Casanada, and D. Steinberg, “Viscosity of culture medium as a regulator of synthesis and secretion of very low density lipoproteins by cultured hepatocytes,” Journal of Biological Chemistry, vol. 257, no. 5, pp. 2188–2192, March 10, 1982 1982.

[39] F. Ye, S. Yin, M. Li, Y. Li, and J. Zhong, “In-vivo full-field measurement of microcirculatory blood flow velocity based on intelligent object identification,” Journal of Biomedical Optics, vol. 25, no. 1, p. 016003, 2020.

[40] H. Stockman, “Lattice Boltzmann Method for Calculating Fluid Flow and Dispersion in Porous and Fractured Media,” in Gas Transport in Porous Media, vol. 20, C. Ho and S. Webb, Eds. (Theory and Applications of Transport in Porous Media: Springer Netherlands, 2006, pp. 221–242.

[41] F. Zhao, B. van Rietbergen, K. Ito, and S. Hofmann, “Flow rates in perfusion bioreactors to maximise mineralisation in bone tissue engineering in vitro,” Journal of Biomechanics, vol. 79, pp. 232-237, 2018/10/05/ 2018.

[42] F. Shen, X. Li, and P. C. H. Li, “Study of flow behaviors on single-cell manipulation and shear stress reduction in microfluidic chips using computational fluid dynamics simulations,” (in eng), Biomicrofluidics, vol. 8, no. 1, pp. 014109–014109, 2014.

[43] A. Wyma, L. Martin-Alarcon, T. Walsh, T. A. Schmidt, I. D. Gates, and M. S. Kallos, “Non-Newtonian rheology in suspension cell cultures significantly impacts bioreactor shear stress quantification,” Biotechnology and Bioengineering, vol. 115, no. 8, pp. 2101–2113, 2018.

[44] V. Sharma, A. Jaishankar, Y.-C. Wang, and G. H. McKinley, “Rheology of globular proteins: apparent yield stress, high shear rate viscosity and interfacial viscoelasticity of bovine serum albumin solutions,” Soft Matter, 10.1039/C0SM01312A vol. 7, no. 11, pp. 5150–5160, 2011.

[45] M. M. Castellanos, J. A. Pathak, and R. H. Colby, “Both protein adsorption and aggregation contribute to shear yielding and viscosity increase in protein solutions,” (in eng), Soft Matter, vol. 10, no. 1, pp. 122–31, Jan 7 2014.

[46] M. Brust, C. Schaefer, L. Pan, M. Garcia, P. Arratia, and C. Wagner, “Rheology of Human Blood Plasma: Viscoelastic Versus Newtonian Behavior,” Physical Review Letters, vol. 110, p. 078305, 02/17 2013.

[47] S. Varchanis, Y. Dimakopoulos, C. Wagner, and J. Tsamopoulos, “How viscoelastic is human blood plasma?,” Soft Matter, 10.1039/C8SM00061A vol. 14, no. 21, pp. 4238–4251, 2018.

[48] C. Graf and J. P. Barras, “Rheological properties of human blood plasma — A comparison of measurements with three different viscometers,” Experientia, vol. 35, no. 2, pp. 224-225, 1979/02/01 1979.

[49] M. A. Tung, “Rheology of Protein Dispersions,” Journal of Texture Studies, vol. 9, no. 1-2, pp. 3–31, 1978.

[50] J. Gonzalez-Molina et al., “Extracellular fluid viscosity enhances liver cancer cell mechanosensing and migration,” Biomaterials, vol. 177, pp. 113-124, 2018/09/01/ 2018.

[51] T. Kitagawa, T. Yamaoka, R. Iwase, and A. Murakami, “Three-dimensional cell seeding and growth in radial-flow perfusion bioreactor for in vitro tissue reconstruction,” Biotechnology and Bioengineering, vol. 93, no. 5, pp. 947–954, 2006.

[52] A. Yazdani, X. Li, and G. Em Karniadakis, “Dynamic and rheological properties of soft biological cell suspensions,” (in eng), Rheologica acta, vol. 55, no. 6, pp. 433–449, 2016.

[53] D. Massai et al., “A Versatile Bioreactor for Dynamic Suspension Cell Culture. Application to the Culture of Cancer Cell Spheroids,” (in eng), PloS one, vol. 11, no. 5, ppp. e0154610-e0154610, 2016.

[54] S. Naskar, V. Kumaran, and B. Basu, “On The Origin of Shear Stress Induced Myogenesis Using PMMA Based Lab-on-Chip,” ACS Biomaterials Science & Engineering, vol. 3, no. 6, pp. 1154-1171, 2017/06/12 2017.

[55] M. G. Park, H. Jang, S.-H. Lee, and C. J. Lee, “Flow Shear Stress Enhances the Proliferative Potential of Cultured Radial Glial Cells Possibly Via an Activation of Mechanosensitive Calcium Channel,” (in eng), Experimental neurobiology, vol. 26, no. 2, pp. 71–81, 2017.

[56] R. Xue and S. Cartmell, “A simple in vitro biomimetic perfusion system for mechanotransduction study,” Science and Technology of Advanced Materials, vol. 21, no. 1, pp. 635-640, 2020/01/31 2020.

[57] S. K. Dash, V. Sharma, R. S. Verma, and S. K. Das, “Low intermittent flow promotes rat mesenchymal stem cell differentiation in logarithmic fluid shear device,” Biomicrofluidics, vol. 14, no. 5, p. 054107, 2020.

